# PhyloRNA: a database of RNA secondary structures with associated phylogenies

**DOI:** 10.64898/2026.03.17.712412

**Authors:** Michela Quadrini, Luca Tesei

**Affiliations:** School of Sciences and Technology, University of Camerino, Via Madonna delle Carceri 7, 62020, Italy

**Keywords:** Phylogeny, Database, RNA Secondary Structure, RNA Abstractions

## Abstract

The ability to access, search, and analyse large collections of RNA molecules together with their secondary structure and evolutionary context is essential for comparative and phylogeny-driven studies. Although RNA secondary structure is known to be more conserved than primary sequence, no existing resource systematically associates individual RNA molecules with curated phylogenetic classifications. Here, we introduce PhyloRNA, a curated meta-database that provides large-scale access to RNA secondary structures collected from public resources or derived from experimentally resolved 3D structures. PhyloRNA allows users to search, select, and download extensive sets of RNA molecules in multiple textual formats, each entry being explicitly linked to phylogenetic annotations derived from five curated taxonomy systems. In addition to taxonomic information, each RNA molecule is accompanied by a rich set of descriptors, including pseudoknot order, genus, and three levels of structural abstraction—Core, Core Plus, and Shape—which facilitate comparative analyses across sets of molecules. PhyloRNA is publicly available at https://bdslab.unicam.it/phylorna/ and is regularly updated to incorporate newly available data and revised taxonomic annotations.

## Introduction

Ribonucleic acid (RNA) plays a central role in gene expression and cellular regulation, acting both as an information carrier and as a functional molecule in key cellular processes. These RNAs adopt complex three-dimensional conformations that are linked to their biological roles (30), and many structural features remain highly conserved across species despite substantial sequence variability (26; 52). RNA secondary structure (60) provides a biologically meaningful and computationally tractable abstraction of RNA 3D conformation, representing canonical Watson–Crick–Franklin base pairs (A–U and G–C) and the non-canonical wobble interaction (G–U).

Secondary structures can be derived from experimentally resolved 3D models deposited in the Protein Data Bank (PDB) (5; 9; 50) using tools such as 3DNA/DSSR (32), RNAView (61), MC-Annotate (24), BPNet (56), Barnaba (7), RNApolis Annotator (59; 65), or FR3D (57; 41), as well as predicted directly from RNA sequences using computational methods (19; 15).

RNA secondary structure is known to be more conserved than primary sequence during evolution, making it a valuable descriptor for phylogenetic and comparative studies. However, linking RNA secondary structures to the taxonomy of their organisms remains challenging, as existing databases do not integrate structural information with curated phylogenetic classifications.

Several taxonomy resources are available, including ENA (63), SILVA (48; 62), LTP (35), NCBI (20), and GTDB (13; 39). SILVA, LTP, and GTDB primarily focus on Bacteria and Archaea and provide genome-based phylogenies, whereas ENA offers a more general-purpose taxonomy. NCBI also maintains curated sequence-based classifications for loci such as ITS and the small and large ribosomal subunits (SSU and LSU) (37; 54). Although these resources are widely used, they differ in taxonomic scope, naming conventions, and rank definitions, and many organisms are not consistently classified across them. As a result, reconstructing or comparing phylogenies across multiple taxonomies—especially for large RNA datasets—requires substantial manual effort and is prone to error.

In the literature, several attempts have been made to create RNA meta-databases of secondary structures, including RNA STRAND (2) and BpRNA (18), which provide predicted or 3D-derived structures. Large integrative resources such as RNAcentral (53) aggregate RNA sequences from multiple databases and provide functional annotations; however, they are primarily sequence-centric and do not systematically associate individual RNA molecules with curated secondary structures and phylogenetic classifications. Similarly, Rfam (38) focuses on RNA families described by covariance models and consensus secondary structures, rather than providing structural representations for individual RNA molecules or linking them to explicit taxonomic classifications, which are conceptually distinct from functional family assignments. The Comparative RNA Web Site (CRW) (11), and its updated version (12; 16), collect predicted secondary structures for specific noncoding RNA families such as ribosomal RNAs, introns, and tRNAs. Specialized resources such as the tmRNA (66) and SPRNA (1) databases store tmRNA and SRP RNA molecules, respectively. While these resources provide valuable and carefully curated structural data, they do not integrate explicit, curated phylogenetic classifications, nor do they support comparative analyses across multiple taxonomy systems. In particular, CRW encodes phylogenetic information implicitly through file-naming conventions that include generic domain- or organelle-level labels (e.g. Bacteria, Eukarya (mitochondrion), chloroplast). However, these labels are not explicitly linked to a well-defined or versioned taxonomic reference database, and the underlying classification system cannot be unambiguously traced to a specific curated taxonomy. As a consequence, phylogenetic information in CRW remains coarse-grained and difficult to exploit systematically for cross-database or multi-taxonomy comparative analyses.

To our knowledge, no existing resource systematically links RNA secondary structures to multiple curated phylogenies while also enabling large-scale, structure-centric queries and downloads. This gap limits comparative and phylogeny-driven studies that require coherent structural and taxonomic information on a large scale. Table 1 summarizes a qualitative comparison between existing RNA secondary-structure databases and the resource introduced in this work. Other tools have been developed to facilitate the use of phylogenetic information in bioinformatics pipelines. For example, NCBI Taxonomist (20; 8) is a command-line utility for retrieving and managing taxonomic classifications from the NCBI Taxonomy database, enabling the resolution of taxon names, taxonomic identifiers, and lineage information. However the tool does not support cross-taxonomy comparison.

**Table 1.**
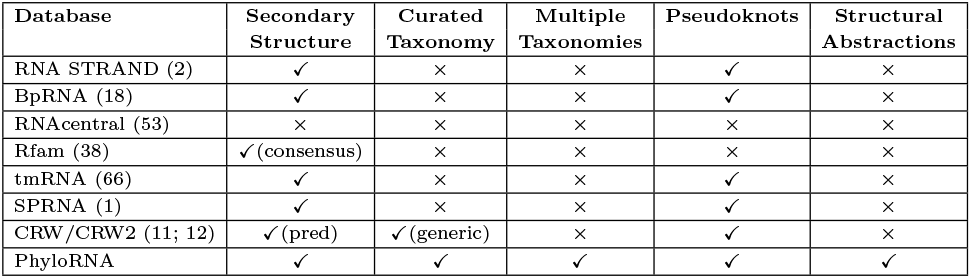
Qualitative comparison between PhyloRNA and existing RNA secondary-structure databases.

In this paper, we introduce PhyloRNA, a meta-database designed to associate RNA secondary structures with curated phylogenetic classifications drawn from the major evolutionary taxonomies. The platform enables users to search for RNA molecules using multiple criteria and to download selected entries in several structural formats, along with their associated metadata, in Comma-Separated Values (CSV) format. Each CSV file includes the taxon labels obtained from the chosen curated taxonomy. Although taxonomic information for individual organisms is readily accessible through public databases, retrieving this information for large sets of RNA molecules is labor-intensive and prone to error when carried out manually. A key advantage of PhyloRNA over existing resources is its ability to associate each RNA structure with taxa from multiple phylogenies automatically and to export extensive collections of molecules in various formats. Switching between taxonomies for downstream analyses requires only changing a selection in the search interface, eliminating the need for repeated manual lookups.

In addition to phylogenetic data, PhyloRNA provides structural descriptors and abstraction levels such as pseudoknot presence and order, genus, and the Shape, Core, and Core Plus abstractions introduced in (44; 45). These features enable analyses that combine evolutionary and structural perspectives, such as identifying conserved motifs or comparing the organization of RNA molecules across taxa. PhyloRNA also incorporates more than 4,000 secondary structures derived from experimentally resolved 3D models in the Protein Data Bank (PDB). These validated structures can be retrieved by selecting “Is Validated = Yes” or filtering entries with “PDB” as their data source. Overall, PhyloRNA provides a unified resource that combines RNA secondary-structure data with curated phylogenetic information, large-scale taxonomy-aware search capabilities, rich structural abstractions, and an expanding collection of experimentally validated PDB-derived structures. The paper is organized as follows. Section 2 describes the technical aspects of the PhyloRNA system. Section 3 presents the web interface, the database content, and the available search criteria. Section 4 illustrates three representative use cases highlighting the advantages of PhyloRNA. Finally, Section 5 summarizes the main contributions and outlines future developments.

## Methods

This section describes the design and implementation of PhyloRNA. We first outline the overall system architecture and the interaction between its main components. We then describe the data acquisition workflow, including the processing of RNA sequences and secondary structures, the computation of structural descriptors, and the integration of curated taxonomic annotations. Finally, we detail the database schema and document organization, as well as the update and validation pipeline used to maintain the resource over time.

### System architecture

PhyloRNA is implemented as a three-tier web application, as illustrated in Figure 1. The current version runs on an Ubuntu Linux virtual machine hosted in our laboratory, which provides all required services.

**Fig. 1.**
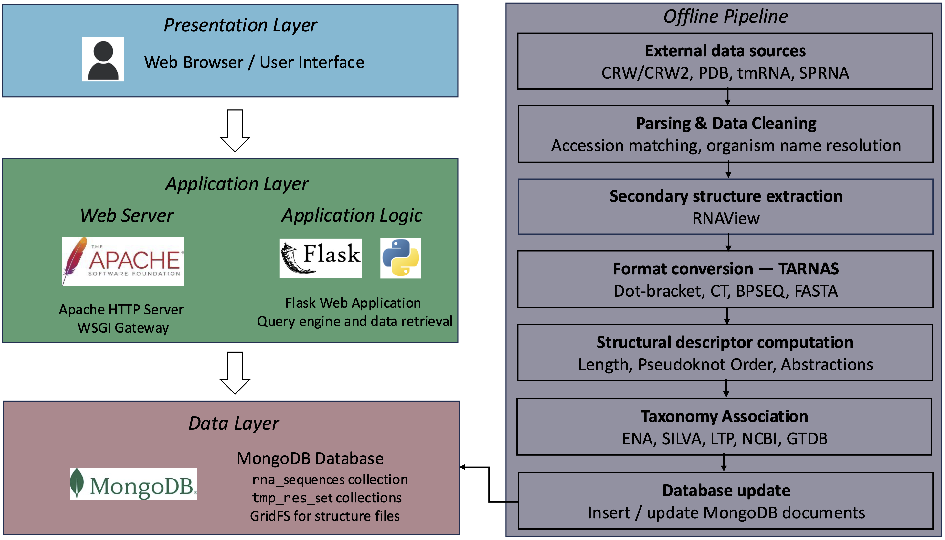
System architecture of PhyloRNA.

The presentation layer consists of a Flask-based web interface accessed through standard web browsers. HTTP requests are handled by an Apache web server configured with a WSGI gateway, which forwards incoming requests to the Flask application.

The application layer implements all search, filtering, and download functionalities. Query parameters submitted by the user are validated and translated into MongoDB queries (34). The data layer is provided by a MongoDB server hosting the main collection `rna_sequences` and temporary query-specific collections `(tmp_res_set)`. RNA secondary-structure files are stored using the GridFS mechanism.

A set of offline scripts and tools orchestrates data acquisition and periodic updates from external resources (CRW/CRW2, PDB, tmRNA, SPRNA, and others). These scripts download the raw data, convert secondary structures into a unified internal representation, compute structural descriptors and abstraction levels, associate each RNA molecule with curated taxonomic classifications from the five supported taxonomy sources, and populate the MongoDB collections.

## Data acquisition workflow

Data are collected from several public resources, including the RCSB Protein Data Bank (PDB) (5; 9; 50), CRW (11), CRW2 (12), the tmRNA database (66), and the SPRNA database (1). For each source, we implemented dedicated parsers that download the original files, extract sequence and structural information, and normalize identifiers such as accession numbers and organism names.

Secondary structures derived from experimentally resolved 3D coordinates are computed using RNAView, whereas predicted secondary structures are taken directly from the original databases. All structures are then converted into a unified internal representation and further translated into multiple output formats using the TARNAS tool. The current release of PhyloRNA provides RNA sequences and secondary structures in the following textual formats:

- **(Extended) Dot-Bracket Notation** (27)
- **BPSEQ** (Base Pairing Sequence) (10)
- **CT** (Connectivity Table) (33)
- **FASTA** (40)

The FASTA format does not encode secondary-structure information but is provided as a sequence-only output option. All formats are stored and exported either with or without header information, according to user selection.

A set of dedicated scripts is used to compute structural descriptors, including sequence length, number of base pairs, number of unpaired nucleotides, pseudoknot order, and genus. Higher-level structural abstractions—namely Shape, Core, and Core Plus—are computed using the TARNAS tool.

Taxonomic annotations are computed for each RNA molecule using Python scripts that query each supported classification system (ENA, SILVA, LTP, NCBI, and GTDB). For each taxonomy, annotations are stored as an array of nested documents that, whenever the organism is classified within a given system, record the complete hierarchy of taxonomic ranks from domain to species.

Finally, all information is assembled into JSON documents and inserted into the MongoDB database.

### Database schema and MongoDB documents

The core collection of PhyloRNA is `rna_sequences`, where each document corresponds to a single RNA molecule and its associated metadata. Each document stores: (i) identifiers such as the internal `reference _id`, the `accession_number`, and the `database_source;` (ii) basic structural descriptors (sequence `length, num_base_pairs`, and `num_unpaired)`; (iii) higher-level structural features, including `is_pseudoknotted, pseudoknot_order, genus,` and the Shape, Core, and Core Plus abstractions; and (iv) curated taxonomic information.

Separate `tmp_res_set` collections store user sessions associated with search and download operations, including query parameters, result identifiers, and selected output formats. Secondary-structure files in multiple formats are stored in GridFS and referenced from the corresponding `rna_sequences` documents. Figure 2 illustrates the UML class diagram of the MongoDB collections.

**Fig. 2.**
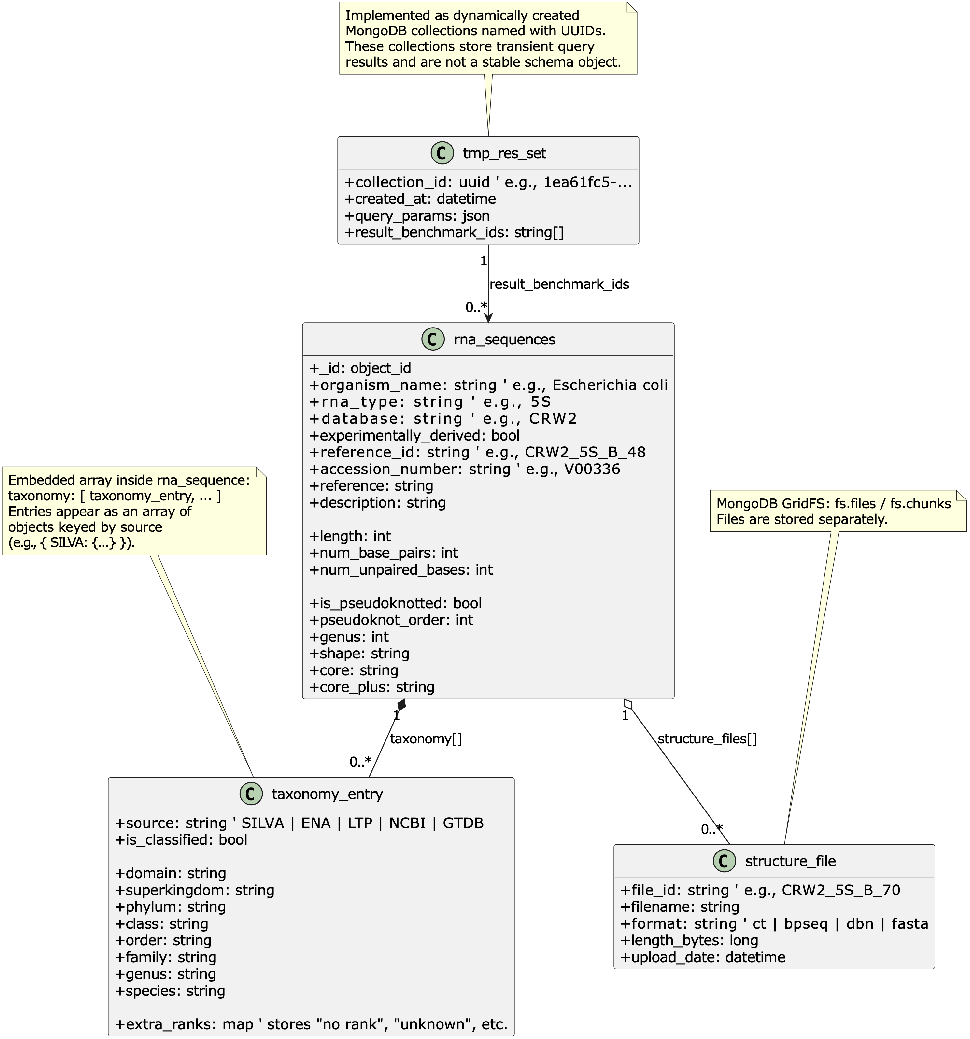
UML class diagram of the MongoDB collections used by PhyloRNA.

### Update and validation pipeline

PhyloRNA is designed to be updated regularly as new RNA structures and taxonomic releases become available. A set of scheduled scripts periodically retrieves the latest versions of the external RNA and taxonomy databases. For RNA secondary structures, the scripts detect newly added or modified entries, process them through the acquisition workflow described in Section 2.2, and insert or update the corresponding MongoDB documents.

Taxonomic annotations are refreshed whenever a new release of SILVA, ENA, LTP, NCBI, or GTDB is published. For each RNA molecule, the updated taxonomy resources are queried and the corresponding taxonomy array is rebuilt.

## Results

This section presents the main features and current content of the PhyloRNA databas (42). The home page displays the number of RNA secondary structures available from each external source, together with their distribution across RNA families, as illustrated in Figure 3.

**Fig. 3.**
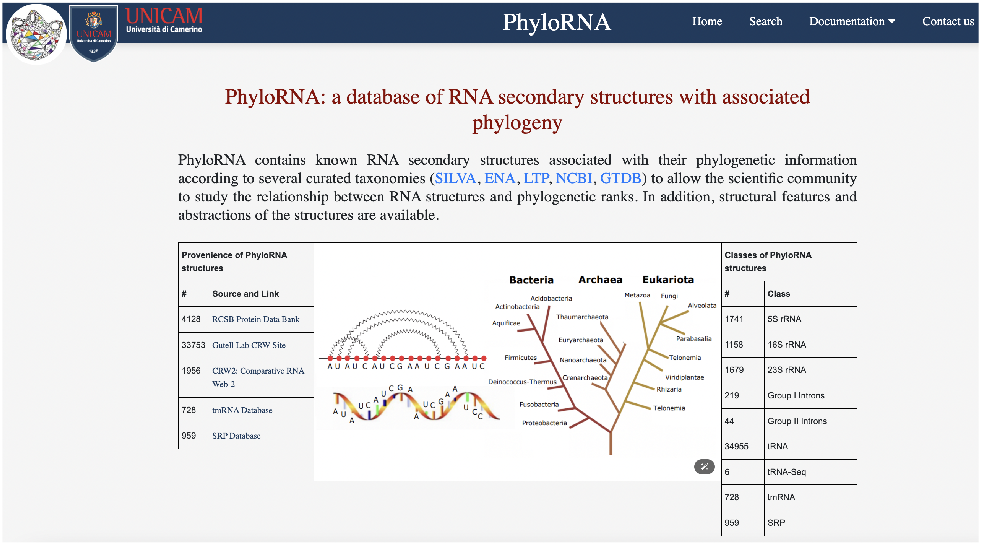
Home page of the PhyloRNA database.

### Search Criteria

PhyloRNA offers several search criteria that users can freely combine to refine their queries. Search criteria can be grouped into identifier-based, size-based, structure-based, and taxonomy-based filters, as described below.

*Identifier* search criteria allow users to search using identifiers associated with RNA molecules.

#### Organism Name

scientific name of the organism associated with the RNA molecule.

#### RNA Type

the family of RNA to which the entry belongs. Examples are ribosomal RNAs 5S, 16S, and 23S; tRNAs; Group I Introns and Group II Introns.

#### Database

the name of the external database from which the entry has been retrieved.

#### Experimentally Derived

a flag indicating whether the RNA secondary structure is derived from an experimentally resolved 3D structure deposited in the Protein Data Bank (PDB). Structures marked as derived are obtained by extracting base-pairing information from experimental 3D models and therefore and therefore correspond to secondary structures derived from experimental 3D data.

#### Reference Id

a unique identifier created for each entry in the PhyloRNA database. Future releases will keep all previous IDs unchanged.

#### Accession Number

identifier of the RNA sequence and its corresponding secondary structure, taken from the external source database.

*Size* criteria **Length, Number of Base Pairs**, and **Number of Unpaired Bases** allow users to specify ranges in order to filter secondary structures by size.

*Structure* criteria rely on the arc-diagram representation of RNA secondary structures. In this representation, nucleotides are placed on a horizontal line and base-pair interactions are drawn as arcs above the sequence, as illustrated in Figure 4.

**Fig. 4.**
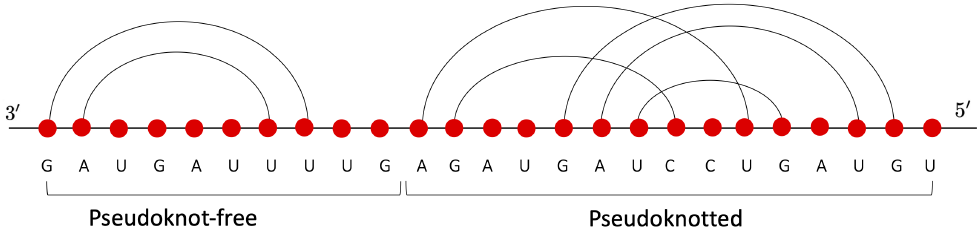
An RNA secondary structure represented by an arc diagram. The base pairs are the zigzagged lines. If arcs in the diagram cross, the structure is pseudoknotted; otherwise, it is pseudoknot-free.

#### Is Pseudoknotted

a flag indicating whether the structure contains at least one pseudoknot. Pseudoknots are widespread RNA motifs involved in several functional roles (17; 58). In an arc diagram, a pseudoknot occurs when two or more arcs cross, introducing challenges for both structure prediction (29) and comparison (47). Figure 4 illustrates an example, with a pseudoknot-free region on the left and a pseudoknotted region on the right.

#### Pseudoknot Order

a measure of the pseudoknot complexity introduced in (4). It is defined as the minimum number of crossing arcs that must be removed from a secondary structure to make it pseudoknot-free. For example, 5S rRNAs are pseudoknot-free and therefore have order 0; 16S rRNAs typically have order 1, meaning that at most two arcs cross; and 23S rRNAs usually fall within orders 2–4, most commonly order 3.

#### Genus

a topological measure of the complexity of an RNA secondary structure. It is defined as the minimum number of handles that must be added to a sphere to obtain a surface on which the arc diagram of the structure can be drawn without crossings. For example, an arc diagram with no crossings can be drawn on a sphere without handles and thus has genus 0. Full details on the definition and computation of genus are provided in (25; 6; 51).

#### **Shape, Core**, and **Core Plus**

abstractions that simplify RNA secondary structures (44; 45). Shapes, introduced by Huang and Reidys (28), are obtained from the arc diagram by removing unpaired nucleotides and non-crossing arcs and collapsing the remaining parallel arcs into one. Figure 5**a** shows the shape corresponding to the molecule illustrated in Figure 4, whose bracket string is ([)].

**Fig. 5.**
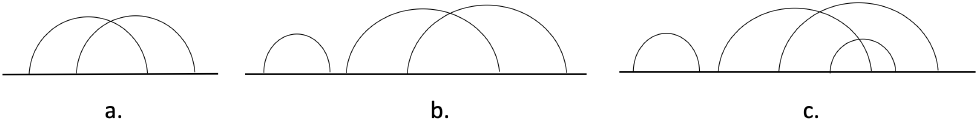
The Shape (**a**), Core (**b**), and Core Plus (**c**) abstractions of the RNA secondary structure depicted in Figure 4.

Core and Core Plus follow similar principles. Core is obtained by first removing unpaired nucleotides and then collapsing all parallel arcs. The core associated with the molecule illustrated in Figure 4 is shown in Figure 5**b**, with bracket string ()([)(]). Core Plus applies these steps in the opposite order—parallel arcs are collapsed first, and unpaired nucleotides are removed afterwards—yielding, for example, the bracket string ()(([)(])), which corresponds to the Core Plus shown in Figure 5**c**.

*Taxonomy* criteria allow users to search RNA molecules based on curated taxonomic classifications and their associated taxonomic ranks.

#### Taxonomy

allows users to select one of the curated taxonomies supported by PhyloRNA, namely ENA (63), SILVA (48; 62), LTP (35), NCBI (20), and GTDB (13; 39).

#### Taxonomic Rank

allows users to select a taxonomic rank within the chosen taxonomy (e.g., superkingdom, phylum, class, order, family, genus).

#### Taxon

specifies the taxon label corresponding to the selected taxonomic rank within the chosen taxonomy (e.g., “Bacteria” as superkingdom, “Proteobacteria” as phylum, “Gammaproteobacteria” as class).

Not all taxonomic systems adopt the same set of taxonomic ranks; consequently, some combinations of taxonomy, rank, and taxon may yield no results. For example, ENA does not use the taxonomic rank `domain`. As a result, querying ENA with the rank `domain` and the taxon “Bacteria” returns no entries. In contrast, performing the same query within GTDB yields all RNA molecules classified as bacteria according to the GTDB taxonomy.

Figure 6 shows an example of search results. Details for each entry can be accessed through the “More Information” column, which also displays the full set of curated taxonomies available for that entry (Figure 7).

**Fig. 6.**
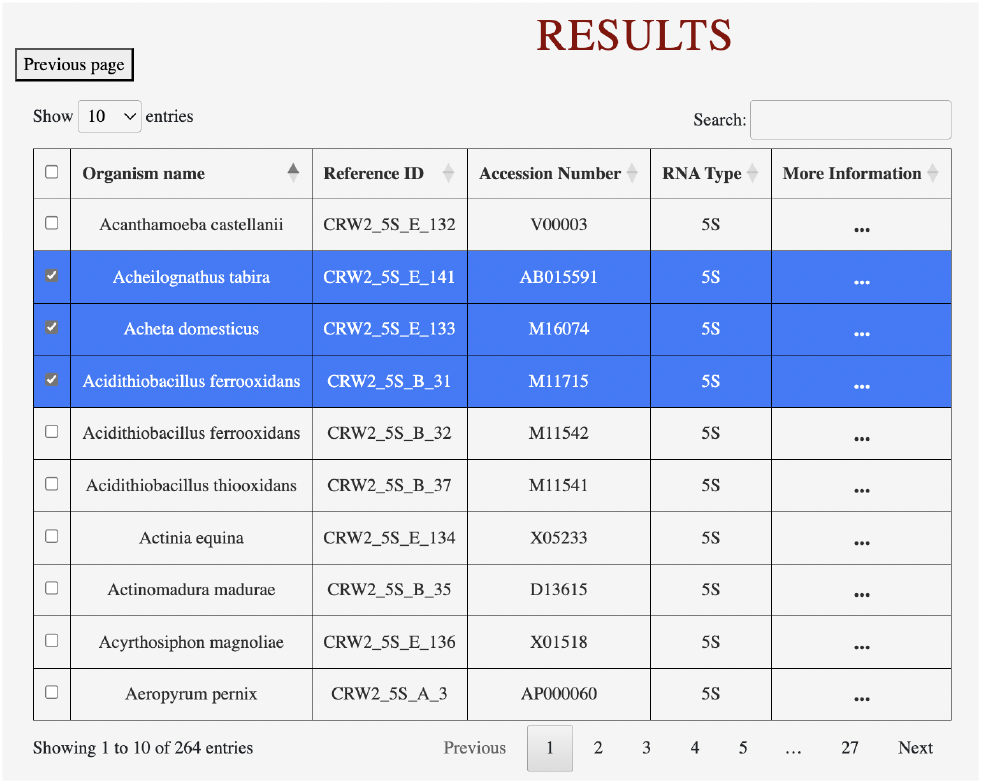
Search results displayed by the PhyloRNA web interface. Each entry can be selected for download. By clicking on “More information” a window is open with the full description of the entry in terms of the search criteria specified in Section 3.1.

**Fig. 7.**
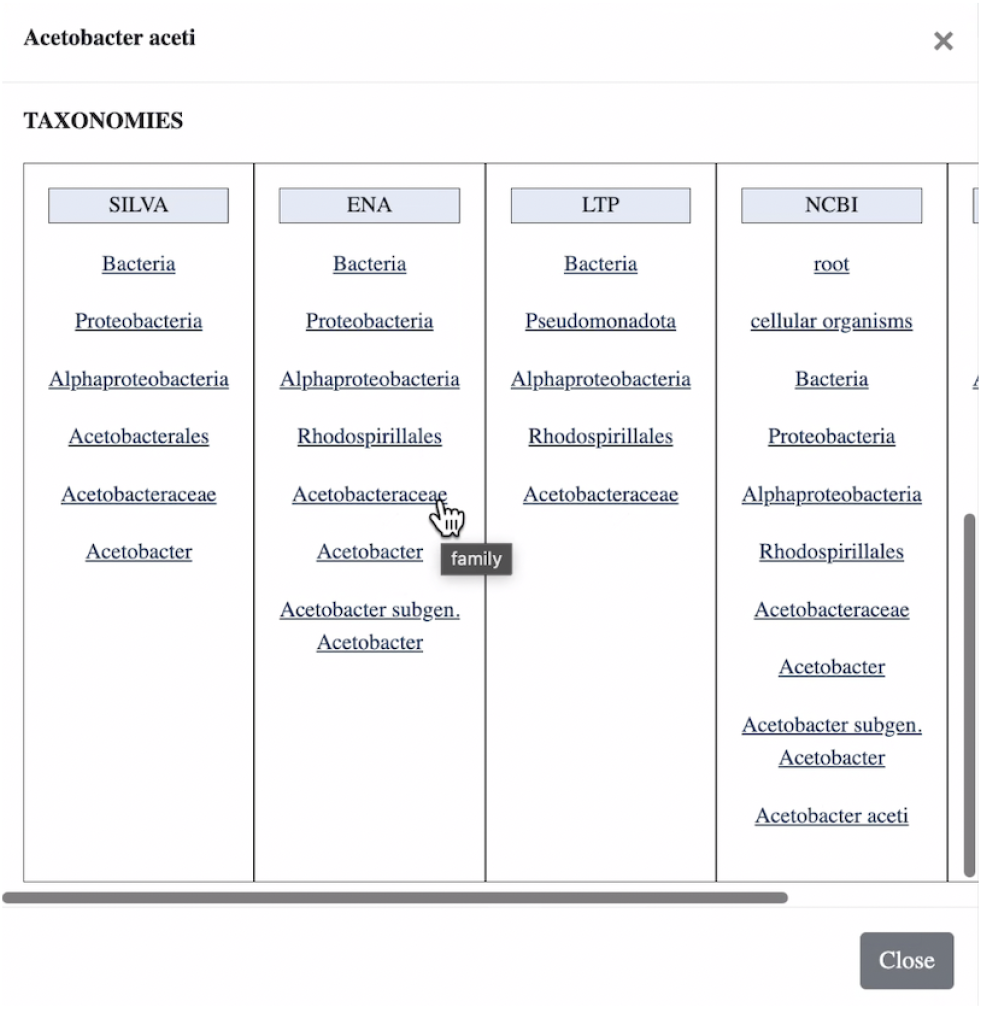
Full description of the phylogenetic information available for a selected entry of results.

### Download

Users can download either the complete set of search results or a selected subset of molecules. The web interface supports pagination with an adjustable number of entries per page and allows users to select individual entries or all entries displayed on the current page. In either case, the corresponding secondary structures can be exported in any of the supported formats (see Section 2.2) and annotated according to the selected curated taxonomy system. All formats can be downloaded with or without header information, which is a useful option when working with tools that do not handle metadata embedded in structure files. Figure 8 shows the interface for selecting the download options.

**Fig. 8.**
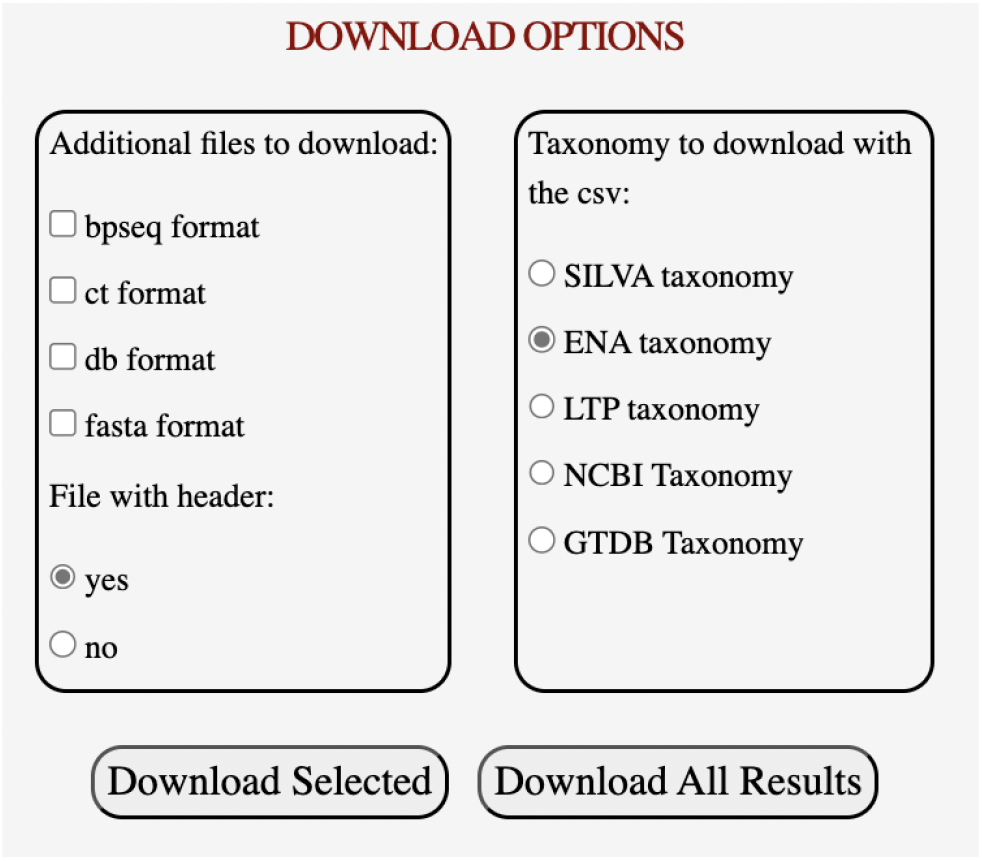
Options to choose the formats of the selected secondary structures to download and the associated phylogeny.

## Applications and Use Cases

The search criteria provided by PhyloRNA—such as taxonomic information, structural features, and abstraction levels— enable the construction of datasets for structure classification, evolutionary analyses, pattern discovery in RNA families, and the evaluation of secondary-structure prediction methods. In this section, we present three use cases that highlight the advantages of using PhyloRNA in these contexts.

### Reconstruction of phylogeny

PhyloRNA provides curated secondary structures and taxonomic annotations that greatly simplify the construction of benchmark datasets for phylogenetic analyses. By enabling users to retrieve large sets of molecules together with consistent taxon labels from multiple curated taxonomies, PhyloRNA removes the need for manual and error-prone data collection, which is often required when reconstructing evolutionary relationships from RNA secondary structures.

This advantage becomes evident when compared with earlier studies. In particular, Quadrini *et al*. (47) introduced a framework for evaluating methods for comparing RNA secondary structures, with a special focus on pseudoknotted RNAs. Their study assessed several distance measures based on Genus(25; 6; 51), PSMAlign (14), ASPRAlign (43; 46), PskOrder (3), and RAG-2D (22; 23; 21; 64). The objective was to determine whether these distances could recover phylum-level taxonomic labels. Constructing the benchmark dataset for this analysis required manually retrieving phylum assignments for Archaea, Bacteria, and Eukaryota 5S, 16S, and 23S rRNAs according to the ENA taxonomy (63), followed by agglomerative clustering.

With PhyloRNA, the entire dataset used in that study could have been assembled automatically, with access to multiple curated taxonomies rather than only ENA. This example illustrates how PhyloRNA supports phylogenetic reconstruction tasks by providing consistent structural and taxonomic information on a large scale.

### Using different taxonomies

PhyloRNA also enables comparative analyses across multiple curated taxonomies, a task that is usually difficult due to the need for manual data collection. To illustrate this, we retrieved all 5S rRNAs from PhyloRNA originating from the CRW2 database. We also downloaded their taxonomic annotations according to each of the five curated taxonomies supported by PhyloRNA. The result is five CSV files containing identical sets of RNA molecules annotated with different taxon labels.

Using these files, we generated five independent classifications by applying hierarchical clustering (36), following the evaluation framework outlined in (47). For simplicity, we used ASPRAlign to compute distances between secondary structures and reconstructed the taxonomy at the superkingdom level (Archaea, Bacteria, Eukaryota). As in (47), clustering performance was assessed using the Rand index, homogeneity, and completeness metrics (49; 55), each ranging from 0 to 1, with higher values indicating better agreement with known labels. We considered the most common linkage strategies: single, complete, and average.

The results, summarized in Table 2, show how clustering performance varies across the five taxonomies. These differences reflect the distinct classification strategies and levels of curation adopted by each taxonomy. Moreover, not all organisms are classified consistently across databases, which further emphasizes the importance of having access to multiple curated taxonomies. PhyloRNA makes such comparative analyses straightforward, enabling researchers to assess how taxonomic choice influences clustering outcomes and to design robust experimental pipelines based on diverse phylogenetic perspectives.

**Table 2.**
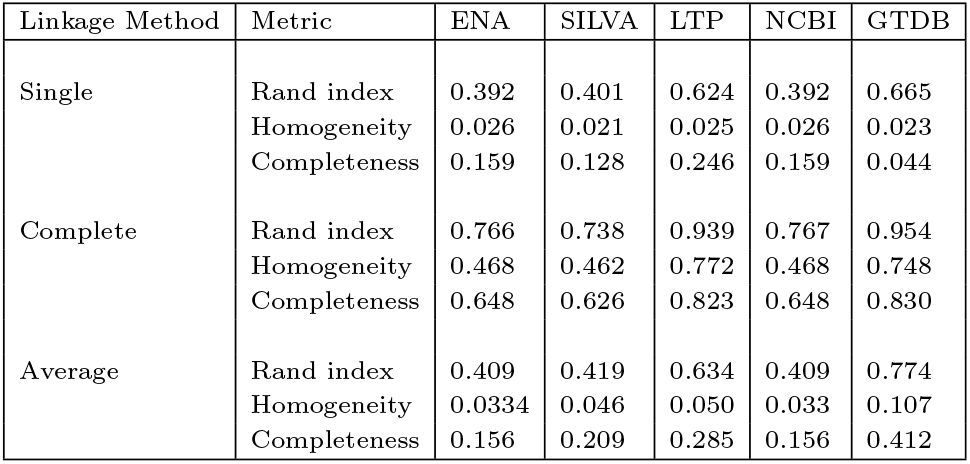
Classification of 5S RNA molecules into the superkingdoms (domain for GTDB) Archaea, Bacteria, and Eukaryota using the ASPRA distance and hierarchical clustering. Clustering was performed using single, complete, and average linkage. Clustering performance was evaluated using the Rand index, homogeneity, and completeness metrics.

### Shape, Core, and Core Plus abstractions

This example illustrates the potential of PhyloRNA to support statistically meaningful analyses that combine RNA structural abstractions with curated phylogenetic classifications. In particular, we focus on the *Shape, Core*, and *Core Plus* abstractions, which reduce the complexity of RNA secondary structures while preserving biologically relevant features (44; 45).

As in the experiment described in Section 4.2, we retrieved all 5S rRNAs originating from the CRW2 database and available in PhyloRNA, and downloaded their taxonomic annotations according to the ENA taxonomy. This dataset comprises 264 RNA secondary structures.

All molecules share the same genus (zero), Shape abstraction (empty, due to the absence of pseudoknots in the considered 5S rRNAs), and Core abstraction ((()())). However, they differ in their Core Plus representations, which are grouped into eight distinct motifs. Among these, two motifs account for the vast majority of the dataset. The Core Plus motif ((((())))((()))) represents 51.5% of the molecules, while ((((())))(())) accounts for 34.8%.

For each of these two dominant motifs, we examined the distribution of ENA superkingdom annotations among the molecules that are classified by ENA. A substantial fraction of molecules lacks an ENA classification and is therefore excluded from the computation of the reported percentages. Specifically, for the first motif, 83 molecules (61%) are not classified by ENA; among the remaining classified molecules, 81.1% belong to the Eukaryota superkingdom. For the second motif, 22 molecules (23.9%) are not classified by ENA; among the classified subset, 94.3% are assigned to the Bacteria superkingdom.

Rather than suggesting a direct structural–phylogenetic correlation, these results demonstrate how PhyloRNA enables the systematic aggregation of structural motifs and curated taxonomic annotations, allowing users to perform large-scale statistical analyses on RNA secondary-structure datasets. The distribution of superkingdoms across the two main Core Plus motifs is summarized in Table 3. Data for the remaining six motifs are not reported, as they account for a negligible fraction of the dataset.

**Table 3.**
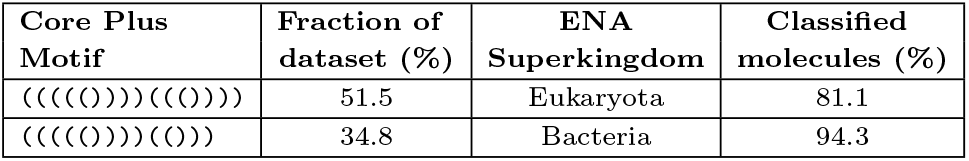
Distribution of ENA superkingdom annotations across the two most frequent Core Plus motifs in the considered dataset of 5S rRNAs. Percentages in the last column are computed only on molecules classified by ENA.

## Conclusion and Future Work

We have presented PhyloRNA, a database of RNA secondary structures enriched with curated phylogenetic classifications. By integrating secondary-structure information with multiple taxonomic systems, PhyloRNA supports a broad range of studies, including phylogenetic reconstruction, comparative analyses of structural features, and the evaluation of RNA secondary-structure prediction and comparison methods. The database also provides access to structural abstraction levels such as Shape, Core, and Core Plus, enabling simplified representations that facilitate large-scale analyses. PhyloRNA is designed to be flexible and extensible, offering a user-friendly web interface that supports complex searches and is freely accessible to the research community.

The current release of PhyloRNA includes 5S rRNA, 16S rRNA, 23S rRNA, Group I Introns, Group II Introns, tRNA, tRNA-seq, tmRNA, and SRP RNA. Future developments will focus on expanding the collection with additional RNA families and with structures derived from other public repositories, including those obtained from experimentally resolved 3D models deposited in the Protein Data Bank (PDB) (5; 9; 50). The database will also be regularly updated with the latest releases of curated taxonomies to ensure consistency with current phylogenetic classifications. Updates of both structural data and taxonomy information are regularly scheduled and reported on the website under “Documentation” → “Sources”.

An important planned extension concerns the integration of a structural pattern–search component into PhyloRNA. This extension will allow users to query the database not only through structural descriptors or taxonomic metadata, but also by specifying precise structural motifs, as proposed in (31). Incorporating this approach into PhyloRNA will further enhance the platform’s analytical capabilities, enabling advanced comparative analyses, motif discovery, and functional annotation.

## Author contributions statement

LT and MQ conceived the idea of the database. MQ collected the relevant initial data. LT was in charge of the technical realization of the database. LT and MQ wrote and revised the paper.

## Acknowledgments

We thank Chiara Cintioni and Denise Falcone for their contribution to the early version of this work. We thank Federico Di Petta for his contribution in fixing the web interface.

This work was supported by the European Union – NextGeneration EU – National Recovery and Resilience Plan (NRRP), Mission 4, Component 2, Investment 1.1, under the PRIN 2022 PNRR call (Min. Decree No. 1409, dated September 14, 2022), project: P2022FFEWN RNA secondary structures and their relationship with function: application to non-coding RNAs (RNA2Fun), CUP: J53D23014960001.

MQ is a member of the GNCS—INdAM.

---

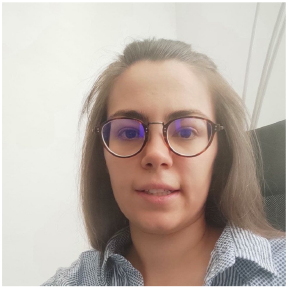

**Michela Quadrini**. received her M.S. degree in Mathematics and her Ph.D. in Computer Science from the University of Camerino, Italy, in 2013 and 2010, respectively. She is currently an Assistant Professor of Computer Science at the University of Camerino. Her research interests include computational systems biology, bioinformatics, artificial intelligence, and formal verification.

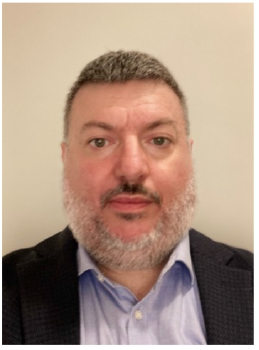

**Luca Tesei** received his M.S. and Ph.D. degrees in Computer Science from the University of Pisa, Italy, in 2000 and 2004, respectively. He is currently an Associate Professor of Computer Science at the University of Camerino, Italy. His research interests include formal verification, bioinformatics, and computational systems biology. He also serves as an Associate Editor for BMC Bioinformatics.

